# Vaccine imprinting drives increased SARS-CoV-2 variant infection in children

**DOI:** 10.64898/2026.08.12.739589

**Authors:** Timothy S. Johnston, Rahul Subramanian, Wakinyan Benhamou, Emily Howerton, Robin Schlesinger, Stephen D. Schmidt, Rachel Kazmierski, Mike Castro, Bob C. Lin, Amy R. Henry, Sarah C. Smith, Jesmine Roberts-Torres, Farida Laboune, I-Ting Teng, Shuishu Wang, Dabeluchi Isiofia, Levi Dong, Mark M. Painter, Alicen B. Spaulding, Chaim A. Schramm, David W. Kimberlin, Samuel R. Dominguez, Hai Nguyen-Tran, Perdita Permaul, E. John Wherry, Matthew R. Vogt, Kevin Messacar, Tongqing Zhou, Leonid A. Serebryannyy, Bryan T. Grenfell, Daniel C. Douek

**Author notes:** Corresponding Author: Daniel C. Douek.

## Abstract

Virus exposure history, particularly first exposure, is believed to shape vaccine efficacy and infection susceptibility; however, evidence for mechanistic links between immune responses in individuals and epidemiological outcome in populations is scarce. Recent co-circulation of SARS-CoV-2 variants XFG and BA.3.2 has revealed a striking enrichment in BA.3.2 cases among children. By combining epidemiological modeling, serology and monoclonal antibody analysis in children and adults, we show the dependence of effective variant-specific antibodies on vaccination history which may explain birth-year influence on differential susceptibility to these co-circulating variants. Ancestral cross-reactive site I antibodies frequently neutralize BA.3.2, but not XFG. By contrast, Omicron type-specific site I/III and III antibodies frequently neutralize XFG but not BA.3.2, revealing a tradeoff in the ability to neutralize these two co-circulating strains. These findings mechanistically link immune history, variant neutralization, antibody repertoire and variant infection risk, and suggest that vaccination regimens in children should prioritize neutralization breadth.

## Main

Decades of studies on influenza virus and, more recently, SARS-CoV-2 infections have suggested that antigen exposure history shapes antibody responses against future, antigenically drifted virus variants^1–9^. At the immunological level, secondary responses to such variants are characterized by the recall of memory B cells and increased titers of cross-reactive antibodies rather than new responses against mutated epitopes, a phenomenon also referred to as imprinting^9–12^. At the epidemiological level, studies evaluating vaccine efficacy and protection against influenza virus variants have revealed disparities associated with birth year and imprinting variant^13–20^. Although they are hypothesized to be related, the immunological and epidemiological outcomes of imprinting have not been bridged.

In North America, the primary vaccine schedule included the ancestral Wuhan-Hu-1 (WT) strain until September 2023, when guidance was updated to the monovalent XBB.1.5 formulation. As such, children born after September 2023 would not have been exposed to ancestral SARS-CoV-2, establishing an Omicron-imprinted “birth-year” cohort. Although the immunological consequences of Wuhan-Hu-1 imprinting in SARS-CoV-2 have been extensively studied^21–23^, the immunological and epidemiological consequences of Omicron vaccine imprinting are less understood.

BA.3.2 was initially detected in South Africa in late 2024 and contains over 70 Spike protein mutations which are thought to be the result of prolonged intra-host evolution^24^. BA.3.2 has since circulated at low overall levels globally yet early reports noted proportional increases in circulation in young age groups compared to JN.1-descendant lineages, predominantly XFG^25^. This proportional enrichment suggests a possible role for birth-year imprinting and variant exposure history in the level of susceptibility to these co-circulating variants. Although virological characterization of BA.3.2 and current JN.1-descendant lineages has been performed^26–28^, serological analyses primarily assess adult sera, which have been found to display similar levels of neutralization against BA.3.2 and JN.1-descendant lineages^29,30^.

Here, we focused on immunological comparisons between adults and children aged 0-4 years with ancestral or Omicron vaccine imprinting, as we hypothesized that vaccine-derived immunity could underlie the difference in variant-specific susceptibility. To this end, we performed a combination of epidemiological modeling, advanced serological analysis and high-throughput monoclonal antibody characterization to determine how immune history shaped clinically relevant antibody responses. We found significant differences in the ability to neutralize virus variants across groups, including BA.3.2 and XFG, dependent on the imprinting vaccine and number of vaccines received. Characterization of variant specific antibodies from these groups uncovered distinct antibody sequences and functions that provide a mechanistic link between the immunological and epidemiological outcomes of imprinting.

### The evolutionary epidemiology of co-circulating XFG and BA.3.2 variants

Recent observations have shown an increased proportion of BA.3.2 cases among young children^24,31^. To observe the differences in variant transmission by age groups, we collected patient status metadata associated with GISAID^32^ SARS-CoV-2 sequences from New York and New Jersey between 2025-01-05 to 2026-07-04 (**Fig. 1a**).

**Figure 1.**
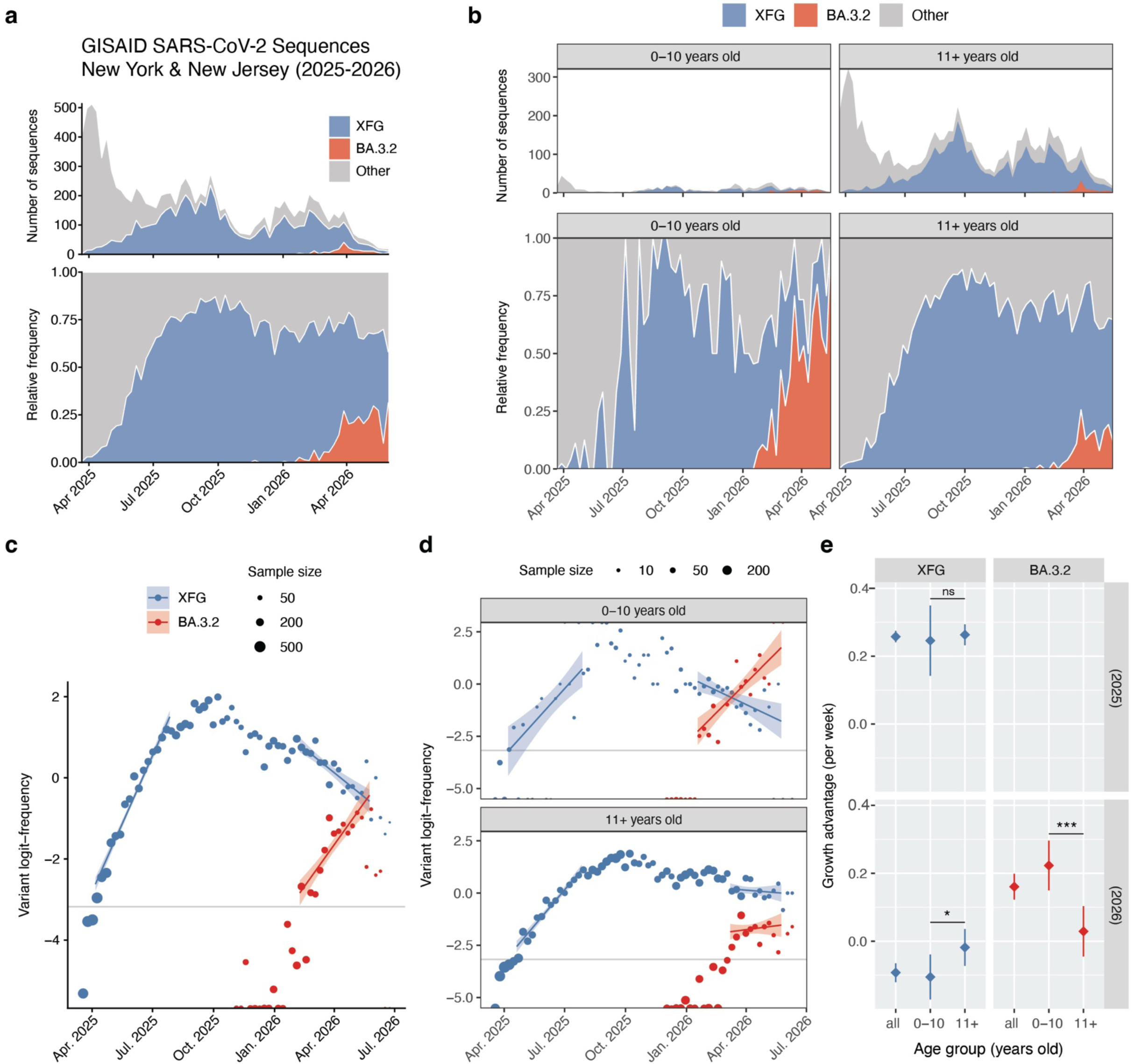
Growth of the XFG and BA.3.2 variants in New York and New Jersey 2025-2026. a) SARS-CoV-2 sequences with patient status metadata from the GISAID database for New York and New Jersey Jan 2025 – Jul 2026, displayed as number and relative frequency. b) As in (a) but stratified by age group. c) XFG and BA.3.2 logit frequency over time, overlaid with logistic regressions. Lines represent model-predicted means, and colored bands the associated 95% confidence intervals. d) As in (c) but stratified by age group. e) Growth advantage estimates per week for XFG (2025 and 2026) and BA.3.2 (2026 only) variants. Diamonds denote mean estimates and whiskers denote 95% confidence intervals. Reported p-values indicate the significance of the age group × time interaction term for each logistic regression model, *p < 0.05, *** p < 0.001.

Stratifying by ages 0-10 years old and 11+ years old revealed BA.3.2 to invade earlier in the younger age group and grow faster, displacing XFG in early 2026 (**Fig. 1b**). During the same period, XFG appeared to remain dominant in the 11+ years old age group.

These data were next used to fit logistic regression models, with the rate of change in variant frequency being estimated on the logit scale as a measure of the relative growth advantage or disadvantage of the variant of interest (BA.3.2 or XFG) compared to all other circulating variants. Three regressions were fitted, one in 2025 for XFG during its emergence, and two in 2026 for XFG and BA.3.2 during the emergence of BA.3.2 (**Fig. 1c**). During its emergence in 2025, XFG appeared to demonstrate a similar growth advantage between age groups. Conversely, during its emergence in 2026, BA.3.2 displayed a significantly higher growth advantage in the 0-10 age group compared to the 11+ age group (and, during the same period, XFG displayed a negative and significantly lower relative growth rate in the 0-10 age group) (**Fig. 1d, e**). Thus, in contrast to XFG which showed no evidence of age-specific differences during its emergence, BA.3.2 demonstrated an age-specific growth advantage among children.

### Ancestral- and Omicron-imprinted children reveal serological effects of imprinting

We hypothesized that this growth advantage of BA.3.2 among children aged 0-10 years might result from imprinting with late Omicron subvariants (XBB.1.5), which might elicit higher neutralizing antibody titers against JN.1-lineage subvariants (XFG) compared to BA.3.2 (**Extended Data Fig. 1**). To measure the ability of sera from ancestral and Omicron-imprinted children to neutralize variants, we performed neutralization assays against a panel of SARS-CoV-2 variants in two groups within our paediatric cohort, one group that received a primary ancestral-based vaccine (N = 54, median age = 2.37 IQR = 1.04, median vaccinations = 2, IQR = 1) and one group that received a primary Omicron-based vaccine (N = 14, median age = 2.06, IQR = 1.21, median vaccinations = 2, IQR = 1) (**Fig. 2a, Supplementary Table 1, Extended Data Fig. 2a, c, d**). Omicron- imprinted children fell into 2 categories (N = 12 XBB.1.5, N = 1 KP.2) (**Extended Data Fig. 2e**). Samples from ancestral-imprinted children were predominantly collected during the periods January – June 2023 and January – June 2024 and samples from Omicron-imprinted children were predominantly collected during the periods January – June 2024 and January – June 2025 (**Extended Data Fig. 2f**). We also measured neutralizing antibody titers from serial monitoring samples collected between 2024 and 2026 in a group of 42 adults, all of whom first received the ancestral vaccine and had multiple Omicron exposures (N = 40, median age = 31, IQR = 12, median vaccinations = 5, IQR = 1.25) (**Fig. 2a, Supplementary Table 1, Extended Data Fig. 2b**).

**Figure 2.**
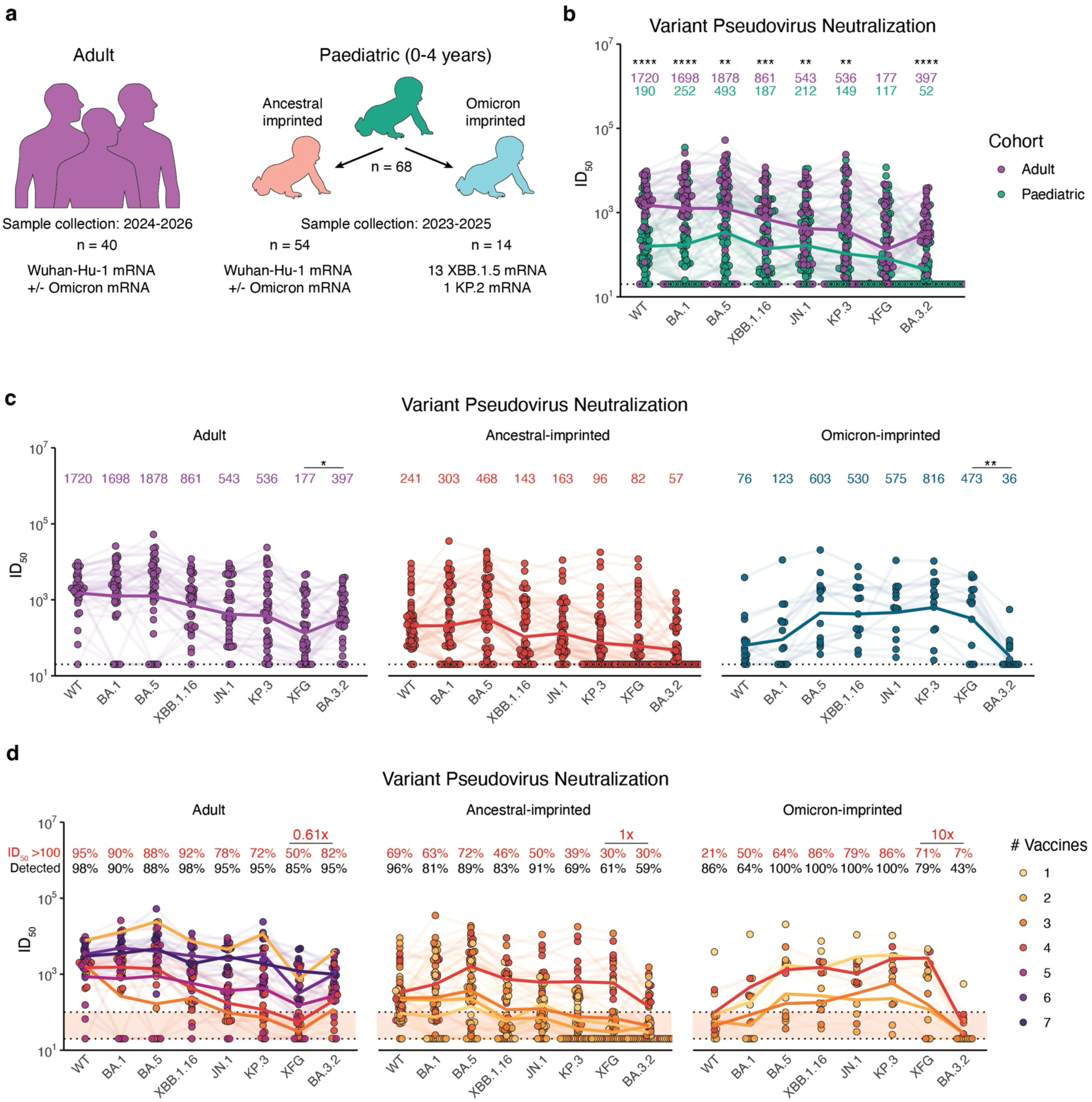
Immune history determines serum neutralization ability. a) Cohort overviews. b) Pseudovirus neutralization inhibitory dilution 50% (ID_50_) values across SARS-CoV-2 variants. Light lines connect individual donor serum; dark lines indicate cohort geometric mean. Geometric mean values reported above, colored by group. c) Pseudovirus neutralization inhibitory dilution 50% (ID_50_) values across SARS-CoV-2 variants. Light lines connect individual donor serum; dark lines indicate cohort geometric mean. Geometric mean values reported above. d) As in c, but lines indicate geometric mean grouped by number of SARS-CoV-2 exposures. In black, percent of serum samples with detectable ID_50_. In red, percent of serum samples above ID_50_ 100. Ratio of neutralization threshold between XFG and BA.3.2 reported above. Dotted lines and shading indicate ID_50_ 20 (LOD) to 100. e) BA.3.2 ID_50_ among individuals with only ancestral exposures, ancestral and Omicron exposures, or only Omicron exposures. Geometric mean reported below. Crossbars represent geometric mean. Wilcoxon rank sum test with Benjamini-Hochberg correction for multiple testing, *p < 0.05, **p < 0.01, ***p < 0.001, ****p < 0.0001.

Adults displayed significantly higher titers than the paediatric cohort across all viruses tested except for XFG (**Fig. 2b**). As we hypothesized that the differences in susceptibility arise from imprinting and not simply age, we further stratified our paediatric cohort into ancestral-imprinted and Omicron-imprinted children (**Fig. 2a**). Neutralization titers varied widely among adults, ancestral-imprinted children, and Omicron-imprinted children. Adults had significantly higher titers against BA.3.2 compared to XFG (**Fig. 2c**). Sera from ancestral-imprinted children followed a similar pattern of neutralization as adults, but at a lower magnitude across all viruses (**Fig. 2c**). Omicron-imprinted children had the highest neutralizing titers against BA.5 through KP.3 and very low titers against WT and BA.3.2. BA.3.2 titers were found to be significantly lower than XFG titers (**Fig. 2c**).

Accounting for the number of vaccines received by each individual, ancestral-imprinted children that received four vaccines had comparable neutralizing titers to adults, including against BA.3.2 (**Fig. 2d**). In the Omicron-imprinted group, neither BA.3.2 titers nor WT titers increased substantially with more vaccine doses (**Fig. 2d**). Across all groups, the total number of vaccines received appeared to associate with greater neutralization titers across variants (**Fig. 2d**). Previous immune correlates analysis of SARS-CoV-2 vaccines has demonstrated that ID_50_ titers greater than 100 correlate with high vaccine efficacy^33–36^. Although it is important to note that our assay is not directly comparable to these studies, we compared the fraction of individuals in each group whose titers were above this threshold for each virus (corresponding to 5x the assay limit of detection), focusing on differences between XFG and BA.3.2. In adults, more individuals reached the BA.3.2 threshold than XFG, resulting in a ratio of 0.61 (**Fig. 2d**). Ancestral-imprinted children had a ratio of 1, denoting equal levels of likely susceptibility (**Fig. 2d**). By contrast, Omicron-imprinted children were above the XFG threshold at a 10-fold greater rate than BA.3.2 (**Fig. 2d**) Together, the higher XFG titers and decreased BA.3.2 titers in Omicron-imprinted children, as well as higher BA.3.2 titers and lower XFG titers in adults provide an immunological explanation for the observed proportional increase in BA.3.2 susceptibility among children aged 0-10 years.

### Antigenic cartography reveals serological niches of BA.3.2 and XFG

Our finding that BA.3.2 neutralizing antibodies are highest in ancestral-imprinted individuals with multiple additional vaccinations led us to hypothesize that variant exposure history is an important determining factor in neutralization breadth. To quantitatively compare the complex serological relationships between sera and virus variants, we constructed an antigenic map^37^ using neutralization titers from primary Wuhan-Hu-1 (N = 8) and primary XBB.1.5 (N = 3) paediatric samples (2x doses mRNA-LNP). Mapping antigenic relationships between SARS-CoV-2 variants in children revealed BA.3.2 to be more antigenically similar to the ancestral strain than XBB.1.16 (**Fig. 3a, Extended Data Fig. 3, Extended Data Table 1**). By contrast, JN.1 and the KP.3/XFG subvariants were more related to XBB.1.16 than the ancestral WT strain (**Fig. 3a, Extended Data Fig. 3, Extended Data Table 2**). These results suggest that initial ancestral exposures benefit BA.3.2 neutralization and initial XBB.1.5 exposures benefit JN.1 subvariant neutralization, including XFG.

**Figure 3.**
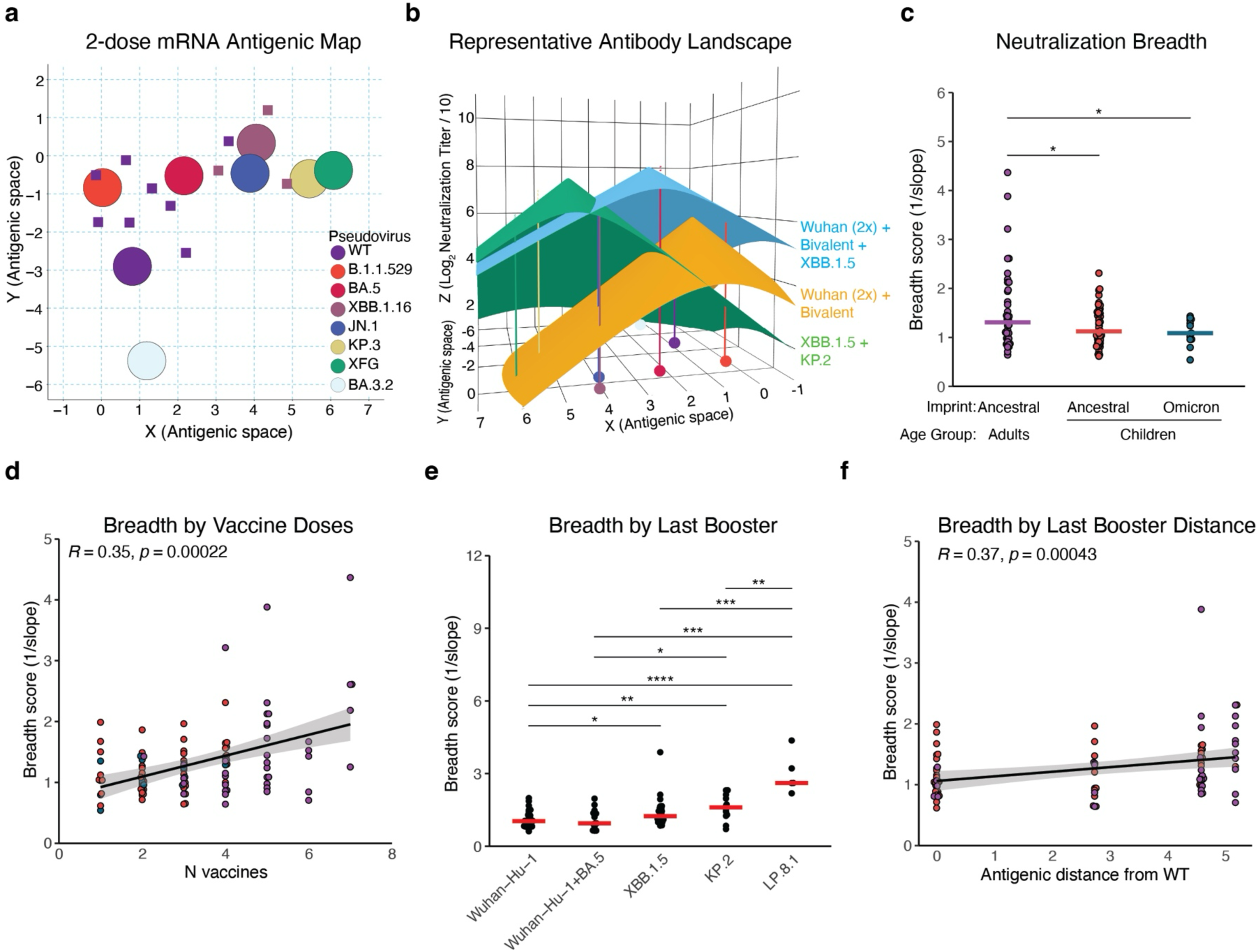
Antigenic cartography and antibody landscapes of adult and child sera. a) Antigenic map constructed from children with 2 doses of an ancestral Wuhan-Hu-1 vaccine (n=9) and children with 2 doses of an XBB.1.5 vaccine (n=2). Circles denote positions of antigen; squares denote positions of sera. Each unit of Euclidean antigenic distance between serum-antigen pairs on the map corresponds to a fold drop in neutralizing antibody titer for that serum against in the antigen in question relative to the antigen against which that person’s serum has the highest neutralizing antibody titer. Sera are colored by their exposure antigen (purple for ancestral, brown for XBB.1.5). b) Representative antibody landscapes, using base map from (a). Group slopes are colored and vaccination history is described on the side. c) Individual antibody breadth scores derived antibody landscape slopes by group. d) Linear regression of antibody breadth score and number of vaccines. Spearman correlation is shown. Shaded band represents pointwise 95% confidence interval. e) Breadth scores grouped by terminal exposure variant. f) Of ancestral-imprinted individuals, linear regression of antibody breadth score and antigenic distance from WT of last booster vaccine received. Spearman correlation is shown. Shaded band represents pointwise 95% confidence interval. Crossbars represent median. For c), multiple t-tests with Benjamini-Hochberg correction. For the rest, Wilcoxon rank sum test with Benjamini-Hochberg correction, *p < 0.05, **p < 0.01, ***p < 0.001, ****p < 0.0001.

To quantify the breadth of serum neutralization, antibody landscapes were generated for each participant across the virus panel, fitting a cone-shaped landscape slope^1,38^. The slope parameter summarizes the rate at which neutralization activity declines across antigenic space. Thus, lower slope values indicate flatter antibody landscapes and broader neutralization, whereas higher slope values indicate more narrow neutralization (**Fig. 3b**). As a representative visualization, we constructed shared-slope average antibody landscapes for three specific exposure histories: individuals who received 2 doses of an ancestral Wuhan-Hu-1 vaccine followed by a bivalent Wuhan-Hu-1/BA.5 vaccine (n=10), individuals who received those same vaccines followed by a subsequent XBB exposure (n=6), and individuals who had two exposures to XBB followed by an exposure to KP.2 (n=2). The patterns in the group slope landscape reflect those observed in the serological data: individuals with both ancestral and late Omicron exposures have high titers against both BA.3.2 and XFG and individuals with only late Omicron subvariant exposures have higher titers against XFG than individuals with only ancestral/early Omicron exposures, but slightly lower titers against BA.3.2 (**Fig. 3b**).

We report the inverse of landscape slope, which we term “breadth score”. In an analysis of antibody landscapes across all individuals, breadth of neutralization was significantly higher in adults than both paediatric groups (**Fig. 3c**). To test our hypothesis that sequential variant exposure increased neutralization breadth, we assessed the relationships between breadth, vaccine number, and exposure sequence using linear models. Increased number of vaccine doses correlated significantly with neutralization breadth (**Fig. 3d**). Within ancestral-imprinted individuals (adults and children), different final variant exposures revealed large differences in neutralization breadth, with individuals only exposed to Wuhan-Hu-1 displaying the lowest breadth and individuals with exposures spanning greater antigenic distances displaying greater breadth (**Fig. 3e, f**). Thus, these serological and cartographic analyses provide immunological support for our epidemiological hypothesis that differing susceptibility to XFG and BA.3.2 is the result of immunological imprinting.

### Association between immune breadth and variant susceptibility

To link our analysis of immune breadth to variant-specific susceptibility, we performed a complementary analysis of titers against BA.3.2 and XFG using a combination of LASSO and linear regressions. First, we performed LASSO regressions for variable selection to predict titers against BA.3.2 and XFG separately, using as predictor variables an individual’s landscape slope, age, column basis titer (maximum titer against any measured antigen), and the antigenic distance between the predicted variant and either their primary or last exposure variant. We then performed linear regressions using all predictors with non-zero coefficients following the LASSO regression.

All examined predictors had non-zero coefficients from the LASSO regression and were thus used in the linear regression models (see **Extended Data Table 3 and 4** for results for BA.3.2 and XFG respectively). For both XFG and BA.3.2, the slope of the cone landscape is a strong predictor neutralization titer, with less steep landscapes (smaller slopes) associated with larger neutralization titers against either variant with large coefficients and p-values < 0.0001. These results support our use of landscape slope as a measure of immune breadth relevant for titers against BA.3.2 or XFG.

We included column basis titer, which is an individual’s highest titer against any of the measured variants, as a proxy to account for individual variation due to high or low responders. A high column basis titer was a significant predictor of neutralization against either variant, with the second-largest coefficients and p-values < 0.0001. This suggests that there is individual variation in immune responses across identical exposure histories that contributes to variant susceptibility.

Larger antigenic distances between the primary exposure variant and the predicted variant were associated with lower titers against BA.3.2 and XFG, although this association is significant for XFG (p = 0.0078) and near reaching significance for BA.3.2 (p = 0.058). These effects are also in line with our epidemiological hypotheses that children with primary ancestral exposures would have imprinted immunity antigenically close to BA.3.2, while children with XBB.1.5 initial exposures would have imprinted immunity antigenically close to XFG.

Smaller antigenic distances between the most recent exposure variant and the predicted variant were associated with increased titers against XFG (p = 0.049) but not BA.3.2 (p=0.151). These results also correspond with our explanation of immune imprinting patterns: whereas boosting with later Omicron variants provides individuals with increased titers against the antigenically close XFG variant, it is unlikely to increase titers against BA.3.2 since late variants are generally more antigenically distant from BA.3.2 than the ancestral strain.

Finally, we found that age is a significant predictor of neutralization titers against both variants (p < 0.0001) although the directionality of the effect varies, with lower age associated with lower titers against BA.3.2 but higher titers against XFG. These results support our epidemiological observations of higher proportional incidence of BA.3.2 in children compared to adults. The higher p-value observed for the distance to the primary variant observed for BA.3.2 could also be due to an interaction between age and exposure history, since all adults have an initial exposure to the ancestral variant while children can have exposures to either the ancestral, XBB.1.5 or KP.2 variants in this analysis. Overall, the results of the statistical model corroborate our serological, epidemiological and cartographic analyses.

Together, these results indicate that individual immune history defines neutralization breadth with priming variant and latest variant exposures being key in defining protection from circulating variants. As it relates to susceptibility to BA.3.2 and XFG, overall breadth, as well as antigenic similarity to the ancestral strain and XBB.1.5 may underlie increased protection in ancestral-imprinted and Omicron-imprinted individuals, respectively.

### Vaccine imprinting profoundly shapes the paediatric B cell repertoire

Few studies have been able to capture differences in B cell repertoire elicited by different primary variant exposures^22,23,39^. To compare the Omicron-specific B cell repertoire from ancestral-imprinted (Wuhan-Hu-1 primary vaccine) and Omicron- imprinted (XBB.1.5 primary vaccine) children and assess whether differences could play a role in variant susceptibility, we sorted XBB.1.5 Spike^+^ and RBD^+^ B cells from 4 donors in each group and performed the Rapid Assembly, Transfection, and Production of Immunoglobulins (RATP-Ig) protocol for the isolation of monoclonal antibodies (mAbs) from single B cells (**Fig. 4a, b, Extended Data Fig. 4a**)^40^. Ancestral imprinted children had higher frequencies of XBB.1.5 Spike-specific and RBD-specific B cells (**Fig. 4c**). From the sorted population, we recovered 451 heavy-light chain pairs, of which 174 were confirmed SARS-CoV-2 Spike reactive after binding and neutralization screening against WA1, XBB.1.16, XFG, and BA.3.2 variants (**Fig. 4d, Extended Data Fig. 4b**).

**Figure 4.**
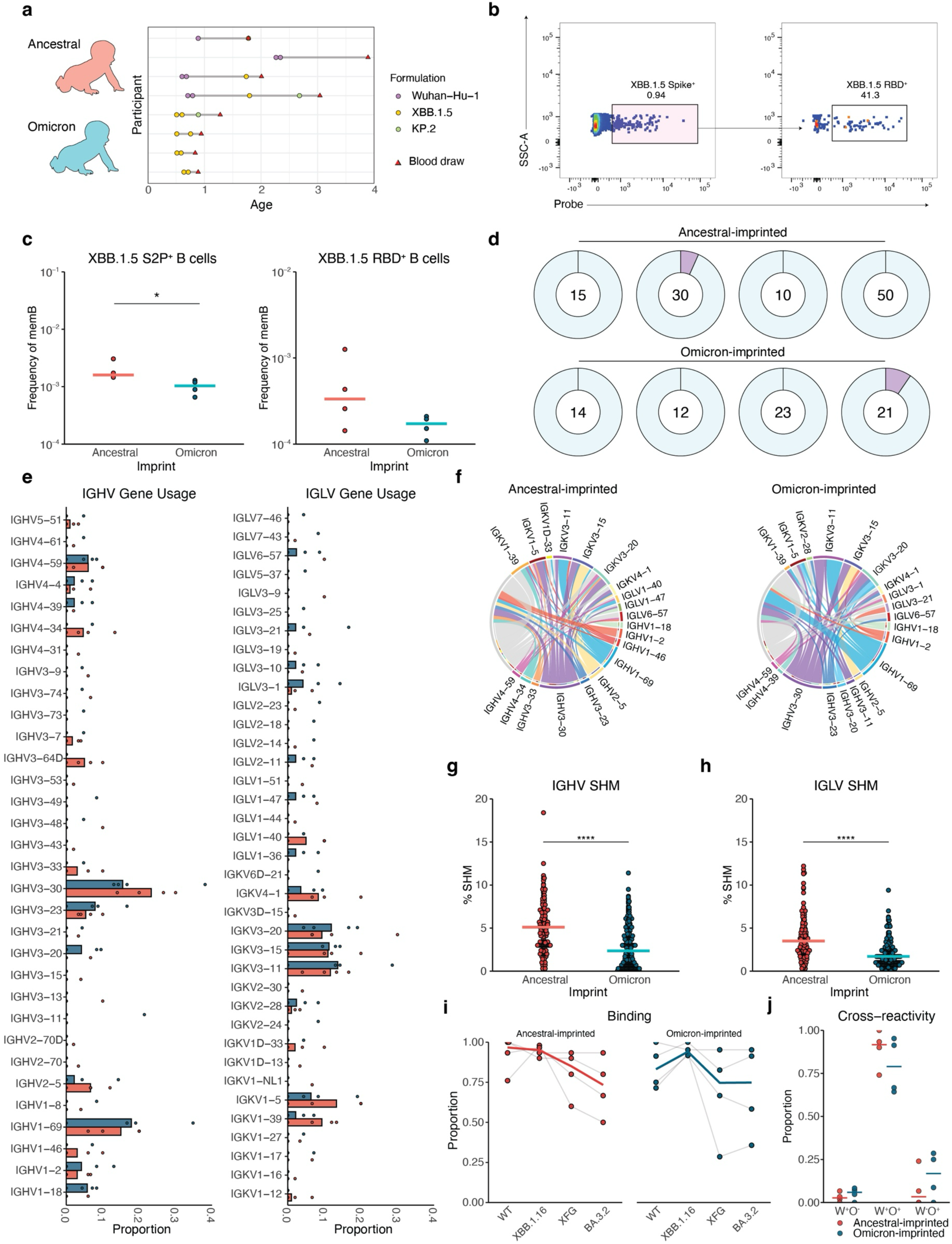
Vaccine imprinting establishes distinct antibody repertoires in children. a) Exposure histories and sampling from paediatric sorting donors. b) Sort gate of XBB.1.5 Spike-specific class-switched memory B cells. c) Frequency of XBB.1.5-Spike (left) and RBD (right) specific B cells among memory B cells. Crossbars represent median. d) Ring plot summary of isolated Spike-specific B cells from each donor. Total N reported in center. Solid light blue indicates singleton proportion, purple segments indicate expanded lineages. e) Proportion of heavy chain (left) and light chain (right) variable gene usage. f) Circos plots of top 10 heavy and light chains with respective pairs per group. Chords connect heavy-light pairings. Inner circle color represents light chain pairing. g) Heavy chain variable gene percent somatic hypermutation. h) Light chain variable gene percent somatic hypermutation. i) Proportion of antibodies binding variant antigens. j) Proportion of antibodies reacting with only WT (W^+^O^-^), WT and Omicron (W^+^O^+^), or only Omicron (W^-^O^+^) antigens. Crossbars represent median. Wilcoxon rank sum test, *p < 0.05, ****p < 0.0001.

We first assessed the frequency of heavy and light chain gene usage between groups (**Fig. 4e**). Genes expressed at higher frequencies in the ancestral-imprinted group included heavy chain variable regions IGHV3-30, 3-64D, 4-34 and light chain variable regions IGKV1-39, 1-5, and 4-1. Genes expressed at higher frequencies in the Omicron-imprinted group included heavy chain variable regions IGHV1-18, 3-20, 4-39, as well as a pattern of increased lambda light chain frequency (**Fig. 4e**).

Notable IGHV:IGLV pairs shared between imprinting groups included IGHV3-30 paired with IGKV1-5, 3-11, 3-15 or 3-20, IGHV1-69:IGKV3-11, IGHV4-39:IGKV1-39, and IGHV2-5:IGKV3-15, suggesting some level of convergence between the ancestral- and XBB.1.5-imprinted repertoires (**Fig. 4f**). We identified pairs found exclusively within each group including IGHV2-5:IGKV3-15, 4-34:IGKV1-39, and IGKV1-15 in the ancestral group and IGHV3-53:IGKV3-20, IGHV4-31:IGLV1-51, IGHV4-39:IGKV3-15, and IGHV4-59:IGLV1-47 in the Omicron group (**Fig. 4f**). Further distinguishing these repertoires was somatic hypermutation of variable genes which was significantly higher in the ancestral group, likely corresponding to increased antigen exposures and cell sorting using a heterologous antigen (**Fig. 4g, h**). Analysis of mAb binding revealed an increase in the proportion of WT cross-reactive antibodies in the ancestral group, and higher frequencies of Omicron type-specific binders in the Omicron group (**Fig. 4i, j**). Collectively, these findings suggest that vaccine imprinting shapes the antibody repertoire at the single B cell level, resulting in changes in the recognition of different SARS-CoV-2 variants.

### Imprinted antibody repertoires determine variant susceptibility

To determine the extent to which the differences in vaccine-elicited repertoires contribute to variant susceptibility, we tested the paediatric antibody panel for neutralization of WT, XBB.1.16, XFG, and BA.3.2 variants (**Extended Data Fig. 5a**). We also assessed neutralization ability of a previously isolated panel of 187 RBD-targeting antibodies that included WT cross-reactive and Omicron type-specific from adults^41^. Antibodies from ancestral-imprinted adults and children neutralized WT virus at a higher frequency than those from Omicron-imprinted children (**Fig. 5a, Extended Data Fig. 5a**). Adults neutralized all variants at greater frequencies, particularly WT, XBB.1.16, and BA.3.2. From the paediatric groups, we isolated 7 XFG-specific nAbs and 2 BA.3.2- specific nAbs. Overall, XFG-specific nAbs also frequently neutralized XBB.1.16, but not WT and BA.3.2. By contrast, BA.3.2-specific nAbs also frequently neutralized WT and XBB.1.16, but not XFG. Few antibodies neutralized both BA.3.2 and XFG, suggesting a tradeoff in the ability to neutralize these two variants (**Extended Data Fig. 5b**). A greater proportion of WT-specific nAbs also neutralized BA.3.2 and a greater proportion of Omicron type-specific (WT non-neutralizing) nAbs neutralized XFG (**Extended Data Fig. 5c**). Thus, WT-reactive antibodies appear to better retain function against BA.3.2 and are evaded by XFG, while the opposite is true for Omicron type-specific antibodies. Adult nAbs were characterized by frequent usage of IGHV1-69:IGLV1-40, IGHV3- 30:IGKV1-39, and IGHV3-53:IGKV1-9 or IGKV1-D33 pairs (**Fig. 5b**). Paediatric nAbs from ancestral and Omicron-imprinted donors revealed distinct pairs, including a predominant IGHV2-5:IGKV3-15 pair, which was not found in adults (**Fig. 5b**). This was the only pair shared between the ancestral and Omicron groups, suggesting divergence in the neutralizing response by priming vaccines.

**Figure 5.**
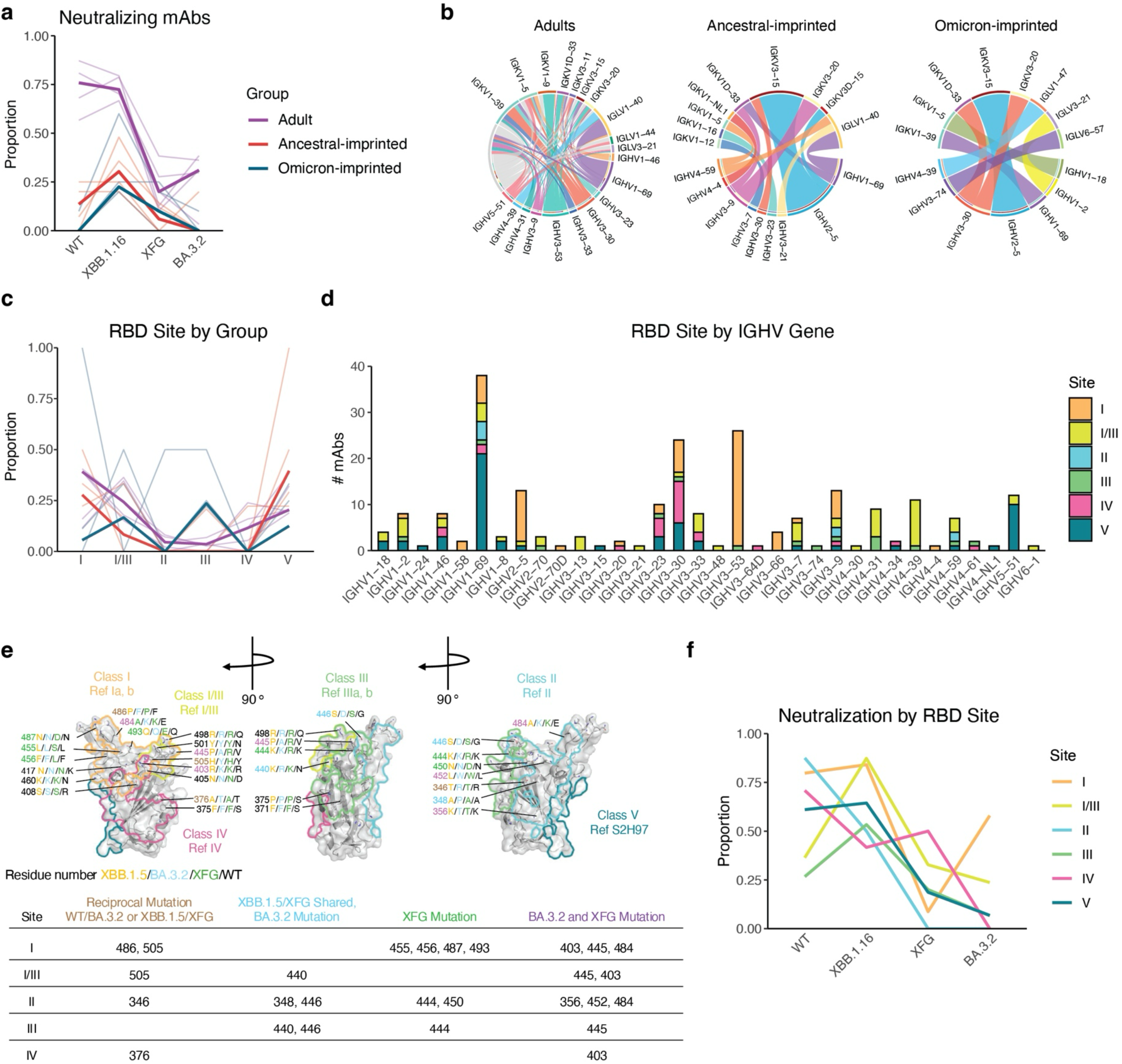
**BA.3.2 escapes Omicron type-specific antibodies in adults and children.**a) Proportion of RBD-targeting antibodies that neutralize WT, XBB.1.16, XFG, and BA.3.2 pseudovirus. Light lines represent individual donor proportions. Dark lines represent group median proportions. b) Circos plots displaying IGHV:IGLV gene pairs of neutralizing antibodies from adults, ancestral-imprinted children, and Omicron-imprinted children. c) Proportion of antibodies targeting each RBD site by group. Light lines represent individual proportions; dark lines represent group median proportions. d) IGHV gene usage of different RBD site targeting mAbs. e) Model of the RBD (PDB:7M7W^44^) with outlined reference antibody footprints. Amino acid substitutions in the epitopes of classes I, I/III, II, III and IV are labeled for WT, XBB.1.5, XFG, and BA.3.2. Residue numbers are colored corresponding to BA.3.2 and XFG mutation. A summary table describing mutations that are mutated in BA.3.2 and XFG at each RBD site is provided. f) Proportion of antibodies targeting each RBD site that neutralize WT, XBB.1.16, XFG, and BA.3.2 variants.

To map RBD binding epitopes, we performed epitope binning with surface plasmon resonance. RBD-binding antibodies of unknown epitope specificity (N = 230) competed against reference antibodies of known specificity for binding to the XBB.1.5 RBD and were binned into binding-site classifications based on their competition profiles. Reference antibodies were previously validated by cryo-EM and cover classes I, II, III, IV, and V^42–44^. In total, we grouped antibodies into 6 major Spike binding sites: I, I/III, II, III, IV, and V (**Extended Data Fig. 5d**).

RBD epitope targeting differed between groups, with more frequent site I, I/III, and V targeting in the adult group, site I and V targeting in the ancestral-imprinted group, and site I/III and III targeting in the Omicron-imprinted group (**Fig. 5c, Extended Data Fig. 5d**). Site I antibodies almost entirely comprised IGHV2-5:IGKV3-15 pairs in paediatric donors and IGHV3-53/66:IGKV1-9 or IGKV1D-33 pairs in adult donors (**Fig. 5d, Extended Data Fig. 5d, e**). BA.3.2 neutralization was largely driven by IGHV3-53/66 lineages (**Extended Data Fig. 5f**). Site I/III comprised many different pairs, with no clear pattern. Site III antibodies had diverse heavy-chain gene usage but were primarily paired with IGLV6-57 or IGKV2-25 (**Fig. 5d, Extended Data Fig. 5e**).

To identify mutations that may mediate variant escape at each neutralizing epitope, we mapped variant mutations onto the RBD structure annotated with the footprints of our reference antibodies (**Fig. 5e**). Most RBD sites had multiple reciprocal mutations, where residues were shared between WT and BA.3.2 or XBB.1.5 and XFG. This included residues 486 and 505 in site I, 505 in site I/III, 346 and 446 in site II, and 376 in site IV.

At these sites, there is therefore a tradeoff in the ability to neutralize XFG and BA.3.2 variants, where XFG evades ancestral-elicited antibodies and BA.3.2 evades late Omicron-elicited antibodies (**Fig. 5e, f**). This also provides explanation for our antigenic cartography findings, where BA.3.2 is more similar to the ancestral strain and XFG is more similar to XBB.1.16 (**Fig. 3a**). Several residues were shared between XBB.1.5 and XFG but mutated in BA.3.2 and WT, including 348, 440 and 446. As such, the BA.3.2 mutations 440R, 446D, and 505Y would contribute to the preferential evasion of Omicron type-specific site I/III and III antibodies (**Fig. 5c, e, f**). Site I had the highest number of XFG mutations at residues that were otherwise shared between WT, XBB.1.5, and BA.3.2, including 455, 456, 487 and 493, explaining the specific escape of ancestral-elicited site I antibodies by XFG (**Fig. 5e, f**). Finally, multiple residues per site were mutated in both XFG and BA.3.2 relative to WT and XBB.1.5, which would evenly contribute to escape from both ancestral-elicited and late Omicron-elicited antibodies.

Together, the mutations found in BA.3.2 and XFG provide an explanation for the site-specific evasion of late Omicron-specific site I/III and III antibodies and ancestral-specific site I antibodies, respectively (**Fig. 5f**). In summary, these data suggest that the antibody repertoires elicited by ancestral- and Omicron-based vaccines demonstrate differences in their ability to neutralize currently circulating variants XFG and BA.3.2 due to divergent RBD epitope targeting.

## Discussion

Here, we demonstrated how immunological imprinting established at the B cell level by vaccination results in both serological and epidemiological consequences. We uncovered a critical gap in immunity within Omicron-imprinted children that results in a tradeoff in the ability to neutralize the co-circulating BA.3.2 and XFG variants. By contrast, ancestral-imprinted adults are at a disadvantage for XFG neutralization, although they maintain greater overall neutralization breadth. Together, these findings reveal how virus variants can take advantage of imprinting-based gaps in antibody immunity to co-circulate.

Mechanistically, we found that ancestral-imprinted individuals have site I antibodies capable of neutralizing BA.3.2, and that BA.3.2 evades the site I/III and III antibodies that are favored by Omicron type-specific neutralizing antibodies. Although children imprinted with Omicron subvariants have higher serum concentrations of neutralizing antibodies against many current variants, our findings suggest that these antibodies target potently neutralizing epitopes that are not shared across diverse strains and thus these children’s overall breadth of immunity is narrower compared to adults.

The important lesson here is that there are circumstances under which an individual, and indeed a population, may require antibodies that target multiple different mutations at the same residue. This suggests that an optimal vaccination approach would be to elicit *de novo* antibodies against that site in previously imprinted people or to immunize with a multivalent vaccine comprising numerous variants at immunodominant sites with high levels of variability in immunologically naïve individuals (i.e. infants).

Between 1980 and 2020, the Yamagata and Victoria lineages of influenza B virus co-circulated, demonstrating preferential circulation dependent on age group and suggesting a birth cohort effect due to immunological imprinting^15^. Our analysis reveals a similar pattern for SARS-CoV-2, as a result of gaps in antibody targeting of variant neutralizing epitopes, whereby more ancestral-related lineages circulate at higher frequencies in Omicron-imprinted individuals and later Omicron subvariant lineages circulate at higher frequencies in ancestral-imprinted individuals. As SARS-CoV-2 lineages continue to diverge, more examples of this may arise.

We further determined that sequential variant exposure confers added breadth, as B cells are recalled if they target “shared” or “conserved” residues across multiple variants. This may better prepare individuals for the sudden emergence of highly divergent variants, such as the case of BA.3.2. We might therefore propose two distinct approaches for conferring neutralization breadth: (1) priming with a multivalent antigen; and (2) sequential vaccination with increasingly antigenically distant antigens. In the former approach, coadministration of multiple antigens would engage multiple pools of B cells specific for diverse virus variants, creating “pockets” of localized neutralization among related viruses. In latter approach, sequential administration would recall and expand memory cells that target shared, possibly potentially conserved, epitopes eliciting a single population of broadly specific B cells. Thus, although both strategies would confer breadth, the B cell repertoires would be distinct.

Childhood vaccination that prioritizes breadth from the outset offers a potential intervention for birth year-linked virus transmission and evolution as observed with influenza virus infection. To confer added breadth, children imprinted with a single Omicron variant may require exposure to the ancestral strain, especially in an immune landscape where it is possible for highly divergent saltation variants to emerge. Furthermore, multivalent vaccines administered as a primary series present a feasible option to include both ancestral and currently circulating Spike variant immunogens in a single formulation. In his 1960 “On the Doctrine of Original Antigenic Sin”, Thomas Francis described the potential benefit of immunological imprinting:

“*Children represent the most susceptible members of the population and probably the most important material for the building of epidemics. The gaps in their immunity should be eliminated by providing early in life the antigenic stimuli to meet the known or anticipated recurrent strains. Natural exposures would then serve to enhance the broad immunity laid down by vaccination. It is our hope that such vaccines can be made from pools of chemically purified antigens-or even with strains experimentally devised. In this manner the original sin of infection could be replaced by an initial blessing of induced immunity*”^5^

There were several limitations worth noting in our experimental design. We have detailed vaccination but not infection records for donors and as such we cannot recreate precise immune histories. Overall, we did not isolate many neutralizing antibodies from children. Indeed, although larger paediatric dataset here would add further resolution, we believe our adult antibody panel provided the clarity required to draw immunological conclusions.

Childhood vaccination rates for SARS-CoV-2 are currently low, with 12.9% coverage for 2024-2025 and 9.7% coverage for 2025-2026 (6 months - 17 years old)^45^. As such, the epidemiological phenomenon we observed could also be attributed to infection-mediated immunity, which was not directly explored here. Still, our main conclusion regarding the contribution of Omicron-imprinted immunity remains.

In conclusion, our data support childhood vaccination regimens that emphasize breadth of immunity, rather than focusing on the most recently circulating variant, to mitigate the immunological constraints established by birth-year imprinting.

## Methods

### Sample Collection

The paediatric cohort here was a subset of donors from the previously described PREMISE clinical research study^46^. The PREMISE clinical research study is a multicentre, prospective, immunoepidemiological surveillance study. Eligible children aged 10 years or younger and weighing at least 8 kg were screened, recruited, and enrolled from inpatient and outpatient settings at three US sites (University of Colorado Anschutz Medical Campus [Aurora, CO, USA], University of North Carolina at Chapel Hill [Chapel Hill, NC, USA], and Weill Cornell Medicine [New York, NY, USA]) in overlapping cohorts from 2022-2025, with the additional enrollment of children at the University of Alabama at Birmingham (Birmingham, AL, USA) from 2023-2024. Briefly, questionnaire data and collection of blood samples (processed to serum, plasma, and peripheral blood mononuclear cells) were collected at two time points: January–June of the year of enrollment (visit 1; pre-enterovirus season), and at follow-up during January–June of the subsequent year (visit 3; post-enterovirus season), separated 6–18 months apart depending on when participants were enrolled and followed up. SARS-CoV-2 vaccine status, including number of doses, vaccine type and date of administration, was collected from vaccination records in the electronic medical record and parent-reported surveys. Samples were selected based on vaccination history and included samples collected during 2023-2024 and 2024-2025. Vaccine immunogen was assumed based on the approved formulation at time of vaccination. This study was approved by the Colorado Multiple Institutional Review Board (COMIRB #21-3488) which served as the sole IRB. Written informed consent was obtained from parents or guardians and assent from participants aged 7 years or older.

Adult subjects were recruited for this study under approval by the University of Pennsylvania IRB (IRB#851465) for blood sampling before and after SARS-CoV-2 mRNA booster vaccination or positive SARS-CoV-2 test as previously described^9,11^. Samples were collected from serial monitoring timepoints between 2024 and 2026. Written informed consent was obtained from all participants. SARS-CoV-2 infection history was determined by self-reporting, and infecting variant was assumed by circulation proportion at time of infection. Participants were otherwise healthy, with no self-reported history of chronic health conditions, and none were hospitalized as a result of a SARS-CoV-2 infection. Venous blood (30–100mL) and clinical questionnaires were collected at each visit.

### Antigenic Cartography and Antibody Landscapes

Antigenic maps were constructed using previously available methods and code from Wang et al.^47^ and Schmidt et al.^48^. To construct our antigenic map, we used samples from children who either received two doses of Wuhan-Hu-1 vaccine or two doses of an XBB.1.5 vaccine. When constructing antigenic maps, we excluded any serum samples with less than 3 detectable antigen measurements from the titer table. We also exclude any children with NP-confirmed infections, defined as having a binding level greater than 7770 AU/mL. We did not use NP-confirmed infections as exclusion criteria when making landscapes, since the landscapes represent a range of heterogenous exposure histories, some of which already have a small sample size. We were more stringent in our selection of samples for the antigen map, since antigenic maps should be constructed using exposure histories of a single antigen (e.g. two doses of an ancestral vaccine). We selected 9 samples from children with two doses of an ancestral Wuhan-Hu-1 vaccine, and 2 samples from children with 2 doses of an XBB.1.5 vaccine.

We assessed the sensitivity of the antigenic map using several approaches including both noisy and resample bootstrapping with 100 replicates per Wang et al. For the noisy bootstrapping, we added random noise with a standard deviation of 0.2878 on a log_2_ scale per the approach of Wang et al. and Schmidt et al. Confidence intervals for the fold drop in titer relative to the ancestral and XBB strains were obtained from the 95 percent quantiles for the noisy bootstrap. We also assessed the dimensionality of the antigenic map by comparing the root-mean square error of maps constructed with 1-5 dimensions (**Extended Data Table 5**).

Landscape fitting and plotting code from previously published antigenic maps and antibody landscapes from Wang et al. and Rossler et al.^49^ were used to construct antibody landscapes. Titers below the limit of detection of 20 were assigned a value of 10 when constructing landscapes. We exclude titer data from seven individuals with unknown exposure histories, as well as from two samples with undetectable titers against all measured antigens.

For all exposure histories, we fit individual cone-shaped landscapes in which each individual has a unique slope and peak location. These slope parameters were used as outcome variables and/or predictors in subsequent statistical models. For representative illustration, we selected three distinct exposure history groups to make average landscapes with a shared slope following the methodology and code of Wang et al. The three exposure groups were individuals who received two doses of a Wuhan-Hu-1 vaccine followed by a bivalent Wuhan-Hu-1/BA.5 vaccine (N = 10), individuals who received two doses of a Wuhan-Hu-1 vaccine, a bivalent Wuhan-Hu-1/BA.5 vaccine, and subsequently an XBB.1.5 vaccine (N = 7), and individuals who received two doses of an XBB.1.5 vaccine followed by a KP.2 exposure (N = 2). The slope of the average landscape is fitted using sera from all samples within that exposure group, with each peak fit separately per individual. To obtain the height of the average landscape, we take the geometric mean of the column basis titer across all samples in the exposure group. For the x- and y-coordinates of the peak of the average landscape, we take the arithmetic mean of the x- and y-coordinates for the peak of each individual landscape in that exposure group.

Cartographic analyses were performed using the antigenic cartography package Racmacs (version 1.2.9, repository https://acorg.github.io/Racmacs/) in RStudio with R version R/4.3.0 and C compiler gcc/11.3.0. Slope plots were constructed using the R package ggplot2 (version 3.5.0), while antibody landscapes were plotted using the R package plotly (version 4.10.3).

### LASSO Regression

We first selected exposure histories for which the last exposure antigen was not on the antigenic map. This excluded participants whose last exposure was to LP.8.1. We then performed Least Absolute Shrinkage and Selection Operator (LASSO) regressions for variable selection to identify predictors of log_2_-transofrmed titers (log_2_(titer/10)) against BA.3.2 or XFG using the R function glmnet() in the R package glmnet, version 4.1-8, following the approach and code of Tshibirani and Hastie (An Introduction to Statistical Learning with R, Version 2, Section 6.5). Optimal lambda values for the regression were selected using the cv.glmnet() function using the entire dataset as the input. We used either raw or standardized individual titers as outcome variables, with the following individual landscape slope, age, column base titer (max titer against measured antigen), and antigenic distance from primary and last exposure antigens as predictors. All predictors had non-zero coefficients for both BA.3.2. and XFG, regardless of whether using log_2_ transformed or standardized titers as outcome variables.

We then fitted linear models of log_2_-transformed BA.3.2 and XFG titers using predictors with non-zero coefficients from the LASSO regression, using the lm() function in the R package stats(). P-values corrected for multiple testing by the Benjamini-Hochberg method were performed using the p.adjust() function in the same package.

### Flow-cytometry Assisted Sorting of Single B cells

Antigen-specific B cells were sorted with biotinylated Spike or RBD streptavidin-fluorophore conjugate probes. All reagents can be found in Supplementary Table 2. Probes were pre-prepared by incubating biotinylated Spike or RBD with fluorescently labeled streptavidin overnight. Each probe included a 5µg:2µg ratio of biotinylated RBD:fluorescently labeled streptavidin. XBB.1.5 Spike and RBD were conjugated to BV605-SA and PECy7-SA, respectively (Biolegend). On the day of sorting, 10e6 PBMCs were thawed from cryopreservation and washed with R10 media (RPMI + 10% FBS + 5% Pen-Strep) + 0.05% Benzonase (Millipore). Cells were stained with 1:500 Ghost Violet 510 Viability Dye (Tonbo) at RT for 10 minutes. Cells were next washed in BSA buffer (BD Biosciences) before surface and probe mastermix was added. Each reaction contained a 50µL surface mastermix of anti-CD3-FITC, anti-CD14-FITC, anti-CD16-FITC, anti-IgD-BB700, anti-CD19-BV421, anti-CD38-BUV661, anti-IgG-Ax700, anti-IgA-PE (final antibody dilution 1:200). A 50µL probe mix of XBB.1.5 RBD-BV605 and XBB.1.5 S2P-PECy7, and 5uM free biotin (Avidity) was added and mixed thoroughly. This reaction was incubated at 4°C for 30 minutes. Cells were then washed, resuspended in 100µL R10, and immediately sorted. All washes were performed by spinning at 500xg for 5 minutes. Single cells were sorted using a BD FACSymphony A5 sorter (BD Biosciences) into 96-well plates with wells containing 5µL Qiagen TCL Buffer (Qiagen) + 1% BME (Sigma-Aldrich) and immediately stored at −80°C. Data were analyzed with FlowJo v10.

### Monoclonal Antibody Isolation and Expression

To isolate heavy and light chain variable regions from single-sorted B cells and express their receptors as monoclonal antibodies, we performed Rapid Assembly, Transfection, and Production of Immunoglobulins (RATP-Ig) as previously described^40^. RNA was extracted from single B cells and 5’ Rapid Amplification of cDNA Ends (RACE) was performed to synthesize and amplify cDNA. Immunoglobulin gene enrichment products were amplified by PCR using primers for IgG, IgA, IgK, and IgL. The enrichment product was assembled into monoclonal antibody DNA expression cassettes using HiFi DNA Assembly Master Mix (New England Biolabs). Heavy and light chain expression cassettes were amplified by PCR, co-transfected into Expi-293 cells (ThermoFisher), and incubated with shaking for 5 days before supernatant collection. Supernatants were clarified by centrifugation at 3000xg for 30 minutes.

### Single Cell V(D)J Sequencing and Analysis

DNA libraries were made from Ig enrichment products using the Illumina Nextera XT library preparation kit (Illumina). The libraries were sequenced on a Nextseq 2000 (Illumina) using a P1 300 cycle kit (Illumina) for paired end 2x150 base pair reads to obtain V(D)J sequences.

Computational analysis of B cell receptor V(D)J sequences was performed as previously reported^40^. Sequencing reads were demultiplexed and V(D)J sequences were assembled using BALDR^50^. Sequences were filtered using a custom script (https://github.com/scharch/filterBALDR) to remove incomplete sequences. Filtered sequences were annotated with SONAR v4.2^51^. The OGRDB database was used to identify and annotate V, D, and J genes using IMGT nomenclature^52,53^. For all analyses, nonproductive rearrangements and cells with multiple heavy or light chains were discarded. All sequences can be found in Supplementary Table 3.

### Total IgG Quantitation

Valita Titer Assay (2.5 - 100 mg/L), 384-well plates (ValitaCell; VAL013) was used according to the manufacturer’s instructions (Beckman Coulter). Undiluted RATP-Ig supernatants and a serially diluted purified IgG reference standard (1-150 µg/mL) were transferred to pre-wetted Valita Titer plates and incubated 15 minutes at room temperature before reading with fluorescence polarization (FP-FLUO detection cartridge, Molecular Devices). Concentrations were interpolated relative to the reference standard.

### Supernatant Binding

Supernatants were screened for binding to the SARS-CoV-2 Spike, RBD, or NTD of WA1, XBB.1.5, XFG, and BA.3.2 by electrochemiluminescence immunoassay using Meso Scale Diagnostic’s (MSD) U-PLEX 96-well plates (K15235N). Biotinylated proteins (200µL at 10 µg/mL) were individually incubated with MSD’s U-PLEX Linkers (Linkers 1- 10 at 300 µL) for 30 minutes at room temperature. The reaction was quenched for 30 minutes with MSD Stop Solution and brought to the manufacturer’s recommended volume for coating. The quenched antigen/U-PLEX linker coating solution was added to plates (50 µL/well), shaking at 900RPM for 1 hour. Plates were washed and supernatants diluted 1:100 in MSD Diluent 100 were added (25 µL/well), shaking at 900RPM for 1 hour. Plates were washed and MSD Sulfo-Tag anti-human IgG detection antibody at 1X (D21ADF; 25µL/well) was added, shaking at 900RPM for 1 hour. Plates were again washed, and MSD Read Buffer B (R60AM; 150µL/well) was added to each well. Plates were read on an MSD S600 Reader. Electrochemiluminescence signals to each respective antigen coated spot in a well was used as a measure of binding. The ECL cutoff for antigen binding was >10,000. MSD values for mAbs can be found in Supplementary Table 3.

### Pseudovirus Neutralization

Serum and mAb supernatant neutralization activity was measured as previously described^54^ using pseudoviruses expressing the spikes of SARS-CoV-2 D614G (referred to as WT), XBB.1.16, XFG, and BA.3.2 variants. Neutralization assays were performed either manually or on an integrated automation platform consisting of a Biomek liquid handler from Beckman Coulter, an ambient temperature labware hotel (Thermo Scientific), a 37°C incubator (Thermo Scientific) and a Molecular Devices Multimode reader. The automated assay methods were operated using Beckman Coulter SAMI EX software (version 5.0). SARS-CoV-2 S-pseudotyped viruses were diluted in 10% FBS, DMEM + 0.3µL/mL puromycin and added into tissue culture plates containing the diluted samples, followed by a 45-minute incubation at 37°C and 5% CO_2_. 293T-human ACE2 reporter cells were added at a concentration of 10,000 cells per well in 20µL into virus/sample tissue culture plates, followed by a 72-h incubation at 37°C and 5% CO_2_. To develop plates, luciferase substrate was added (Revvity Britelite Plus, 6066769). Luminescence signal (relative luminescence unit (RLU)) was measured using a i3x Multimode reader. Percent neutralization of the sample was determined by normalization of the test sample relative light units (RLU) to those of virus and cell control wells with the following calculation: percentage = [(test wells – average of cell control wells) – (average of virus control wells – average cell control wells)] ÷ (average virus control wells – average cell control wells) × 100. For supernatants, neutralization was calculated as % reduction at the dilution tested with a positivity cutoff of >50%. Neutralization values for mAbs can be found in Supplementary Table 3.

### High-throughput Surface Plasmon Resonance

Antibody references of known binding specificity were diluted to 10ug/mL in 10mM sodium acetate pH4.5 solution (Carterra) and amine coupled to an HC30M chip (Carterra) using 100mM Sulfo-NHS and 400nM EDC (Thermofisher). 1M ethanolamine was used to block free polycarboxylate and the chip was washed with 10mM MES + 150mM NaCl + 0.05% Tween 20 buffer (Carterra). Antibody supernatants were diluted to 10ug/mL in HBST (Carterra) + 0.05mg/mL BSA. Epitope binning cycles included 1 minute baseline, 50nM XBB.1.5 RBD injection with 5-minute association time, mAb injection for 5-minute association time, followed by a 1-minute dissociation time, and 30s x 2 regeneration with 10mM Glycine pH 2.0 (Carterra). RU measurements were recorded at 25°C. Analysis was performed using Carterra Epitope Software v.1.9.2.4505 (Carterra). Binary competition measurements were exported, and epitope determinations were made by clustering mAbs with references using Ward’s D2 method. mAbs were assigned manually if clustering assignments conflicted with epitope-defining interactions. Epitope determinations can be found in Supplementary Table 3.

### Structural Visualization

The footprints of the reference antibodies were mapped onto the structure of RBD (PDB:7M7W^44^) using PyMOL version 3.1.6.1^55^. Consensus variant spike protein sequences were downloaded from CoV-Spectrum^56^ and aligned using Clustal Omega^57^.

### Statistical Analysis

All statistical analysis was performed in R (version 4.5.1) unless specifically indicated otherwise in the methods. Shapiro-Wilk testing was performed to test for the normality of data distributions. Non-parametric statistics were used throughout this manuscript unless specifically stated. T-test, Wilcoxon Rank-Sum, Wilcoxon Signed-Rank, and Spearman Rank correlation usage is indicated in figure legends. All tests performed were 2-sided. Crossbars represent median, mean, or geometric mean as indicated in figure legends. Corrections for multiple testing are described in figure legends.

### Data Availability

Data are available upon request from the corresponding author. Supplementary data are available for this paper. Sequencing data have been deposited in Gene Expression Omnibus (GEO) repository and under accession number [pending].

### Code Availability

Code used to construct antigenic maps and antibody landscapes and perform LASSO/linear regressions of BA.3.2 and XFG titers will be made available upon publication in the following GitHub repository: https://github.com/niaid/Manuscript_Repo_EVD_68_COVEND_BA_3_2_Imprinting_Carto R scripts used to create figures are available upon request.

## Acknowledgements

We would like to thank the study participants for making the study possible. We thank the Immune Health team and PREMISE EVD-68 study team for sample collection and processing. We thank David Ambrozak and Daniela Gutierrez for assistance with sorting. We thank Manjula Basappa, Yogita Jethmalani, and Lu (Jennifer) Wang for their assistance with neutralization assays. We thank the VRC VIP Sample Management Team for sample management. We thank Lingshu Wang and Nicole Doria-Rose for SPR reference antibodies.

## Funding

This research was supported in part by the Intramural Research Program of the National Institutes of Health (NIH). The contributions of the NIH author(s) are considered Works of the United States Government. The findings and conclusions presented in this paper are those of the author(s) and do not necessarily reflect the views of the NIH or the U.S. Department of Health and Human Services. This project has been funded in part with federal funds from the National Institute of Allergy and Infectious Diseases, National Institutes of Health, Department of Health and Human Services, under contract no. 75N93021C00015 (E.J.W.) and grant numbers U19AI082630 (E.J.W.), AI105343 (E.J.W.), AI108545 (E.J.W.), AI155577 (E.J.W.), AI149680 (E.J.W.), AI156125 (M.R.V.); and funding from the Parker Institute for Cancer Immunotherapy (to E.J.W.). E.H., D.W.K., S.R.D., H.N.T., P.P., M.R.V., K.M. and B.T.G. received funding through NCI/NIH Contract No. 75N91019D00024, Task Order No. 75N91022F00005 and Task Order No. 75N91023F00016. W.B. receives support from Princeton Catalysis Initiative and the High Meadows Environmental Institute at Princeton University.

## Author Contributions

T.S.J., R.S., L.A.S. and D.C.D. conceived and designed the study. T.S.J., R.S., R.S., S.D.S., R.K., M.C., B.C.L., A.R.H., S.C.S., J.R.T., F.L., L.D., I.T.T. and L.A.S. performed experiments. T.S.J., R.S., E.H., W.B., S.C.S., D.I. and C.A.S. performed bioinformatic analysis. T.S.J., R.S., E.H., W.B., S.W. and D.I. visualized data. M.M.P., A.B.S., H.N.T., S.R.D., K.M., M.R.V., P.P., D.W.K. and E.J.W developed sampling protocols, enrolled subjects, and collected samples/data. T.S.J. wrote the manuscript and all authors provided feedback. T.Z., E.J.W., B.T.G. and D.C.D. provided funding and experimental oversight.

## Competing Interests

E.J.W. advises Arpelos Bioscience, Arsenal Biosciences, Coherus, Danger Bio, IpiNovyx, New Limit, Marengo, Pluto Immunotherapeutics Related Sciences, Santa Ana Bio, and Synthekine. E.J.W. is a founder of and holds shares of Coherus, Danger Bio, and Arsenal Biosciences. All other authors declare no competing interests.

## Corresponding Author

Correspondence and requests for materials should be addressed to Daniel C. Douek.

## Extended Data Figures and Tables

**Extended Data Table 1:** Fold drop in titer relative to the ancestral variant (WT) with confidence intervals obtained from the 95 percent quantiles of the noisy bootstrap replicates.

| Antigen | Fold Drop Relative to WT |
| --- | --- |
| WT | 1 (1 - 1) |
| B.1.1.529 | 4.4 (2.3 - 6.1) |
| BA.5 | 7 (5.7 - 8.2) |
| XBB.1.16 | 24.6 (19.9 - 30.2) |
| JN.1 | 14.9 (11.7 - 24) |
| KP.3 | 37.6 (28.2 - 48.9) |
| XFG | 57.9 (44.5 - 79.7) |
| BA.3.2 | 5.9 (4.6 - 8.1) |

**Extended Data Table 2:** Fold drop in titer relative to XBB.1.16 with confidence intervals obtained from the 95 percent quantiles of the noisy bootstrap replicates.

| Antigen | Fold Drop Relative to XBB.1.16 |
| --- | --- |
| WT | 24.6 (19.9 - 30.2) |
| B.1.1.529 | 17.9 (14.7 - 24.6) |
| BA.5 | 4.2 (3.5 - 5.1) |
| XBB.1.16 | 1 (1 - 1) |
| JN.1 | 1.8 (1.3 - 2.9) |
| KP.3 | 3.3 (2.5 - 4.5) |
| XFG | 4.6 (3.6 - 5.9) |
| BA.3.2 | 86.6 (63.4 - 125.2) |

**Extended Data Table 3:** Results from linear regression model of log_2_-transformed neutralization titers against BA.3.2 as a function of all predictors with non-zero coefficients following LASSO regression. The right-hand column shows p-values corrected for multiple testing by the Benjamini-Hochberg method.

| Variable | Coefficient | Std Error | BH P-value |
| --- | --- | --- | --- |
| (Intercept) | 0.167 | 0.566 | 0.776 |
| Ant_Dist_From_Last_Exposure_Variant | -0.125 | 0.081 | 0.151 |
| Ant_Dist_From_Primary_Exposure_Variant | -0.189 | 0.089 | 0.0579 |
| age | 0.039 | 0.007 | <0.0001 |
| slope | -3.134 | 0.441 | <0.0001 |
| Col_Base_Titer | 0.858 | 0.057 | <0.0001 |

**Extended Data Table 4:** Results from linear regression model of log_2_-transformed neutralization titers against XFG as a function of all predictors with non-zero coefficients following LASSO regression. The right-hand column shows p-values corrected for multiple testing by the Benjamini-Hochberg method.

| Variable | Coefficient | Std Error | BH P-value |
| --- | --- | --- | --- |
| (Intercept) | 3.676 | 0.976 | 0.000482 |
| Ant_Dist_From_Last_Exposure_Variant | -0.232 | 0.116 | 0.0489 |
| Ant_Dist_From_Primary_Exposure_Variant | -0.394 | 0.141 | 0.00783 |
| age | -0.050 | 0.011 | <0.0001 |
| slope | -3.973 | 0.662 | <0.0001 |
| Col_Base_Titer | 0.931 | 0.087 | <0.0001 |

**Extended Data Table 5:** Results from dimensionality testing of antigenic map. We determined the optimal number of dimensions via cross validation with a test set of 100 repeats that each excluded 10% of observed titers. Lower root-mean square errors for detectable titers (above the limit of detection) and undetectable titers (below the limit of detection) are an indicator of the optimal dimensionality of the map.

| Dimensions | mean_rmse<br>detectable | var_rmse<br>detectable | mean_rmse<br>nondetectable | var_rmse<br>nondetectable |
| --- | --- | --- | --- | --- |
| 1 | 1.614 | 0.299 | 1.471 | 1.426 |
| 2 | 1.545 | 0.360 | 1.265 | 0.882 |
| 3 | 1.569 | 0.364 | 1.301 | 0.890 |
| 4 | 1.568 | 0.363 | 1.305 | 0.897 |
| 5 | 1.568 | 0.363 | 1.305 | 0.897 |

**Extended Data Figure 1.**
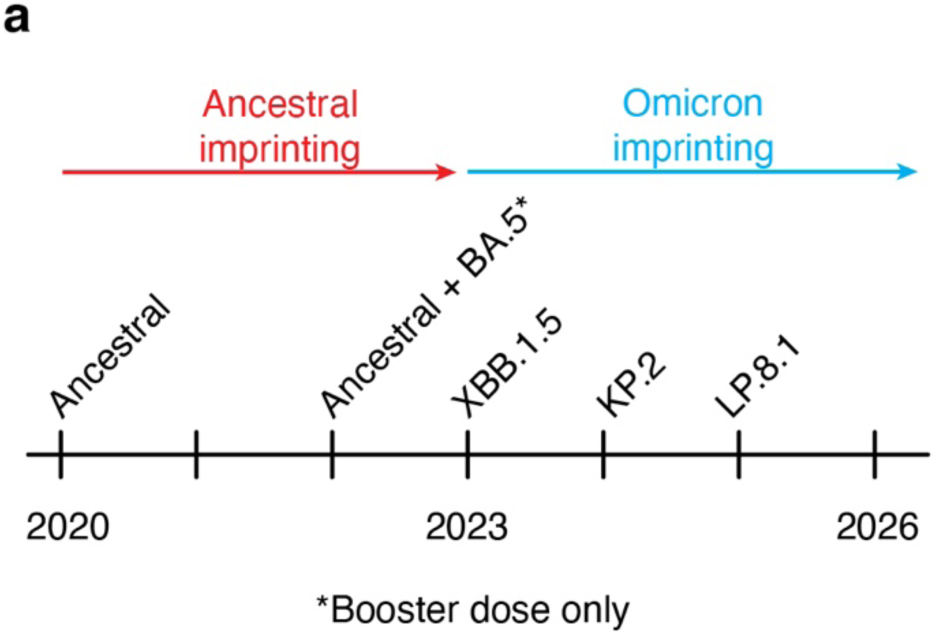
Timeline of vaccine antigens in the United States. a) Timeline of vaccine antigens in the United States. Colored arrows show time periods of vaccine imprinting.

**Extended Data Figure 2.**
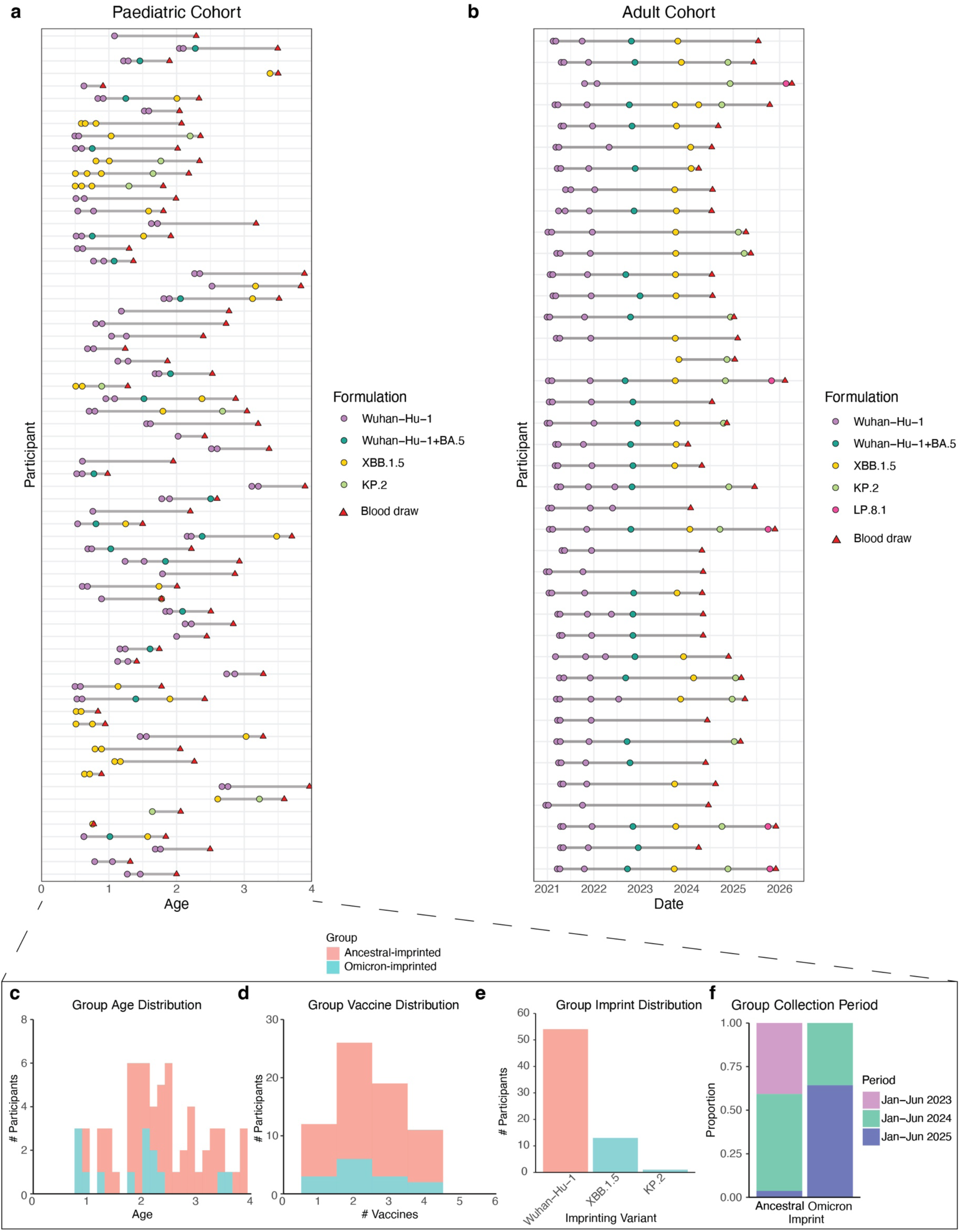
Cohort details. a) Paediatric cohort vaccination and blood draws by age of event. b) Adult cohort vaccination and blood draw by date. Lines connect individual donors. Dots represent vaccines, triangle denotes blood draws. c) Group age distribution of ancestral and Omicron-imprinted children. d) Group vaccine dose number distribution of ancestral and Omicron-imprinted children. e) Distribution of first vaccine variant. f) Proportion of samples collected during each period by group.

**Extended Data Figure 3.**
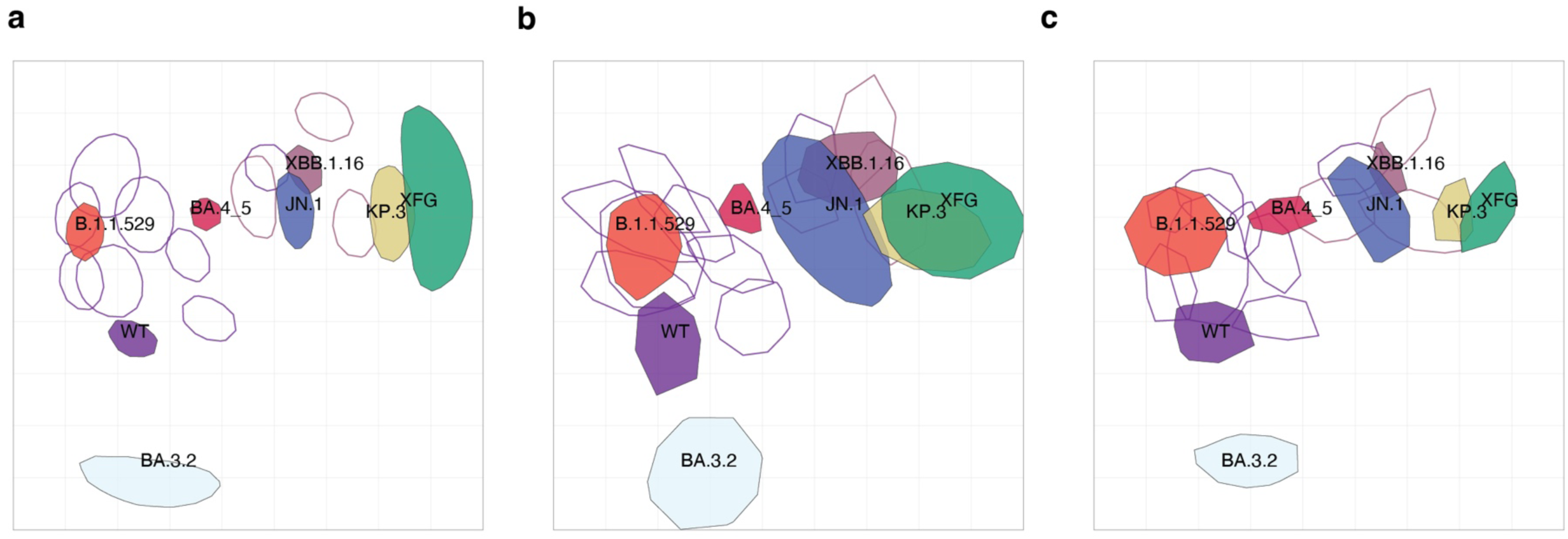
Diagnostic plots for antigenic map for children with 2 doses of Wuhan-Hu-1 or 2 doses of XBB.1.5 vaccination. a) Triangulation confidence intervals. Each shape denotes the area that a single serum or antigen can occupy without increasing the map stress beyond 1 unit of antigenic distance. b) Noisy bootstrap map. An antigenic map is constructed for each of 100 bootstrap replicates with measurement noise added, and the antigen shapes represent possible locations across all replicate maps. c) Resampled bootstrap maps.

**Extended Data Figure 4.**
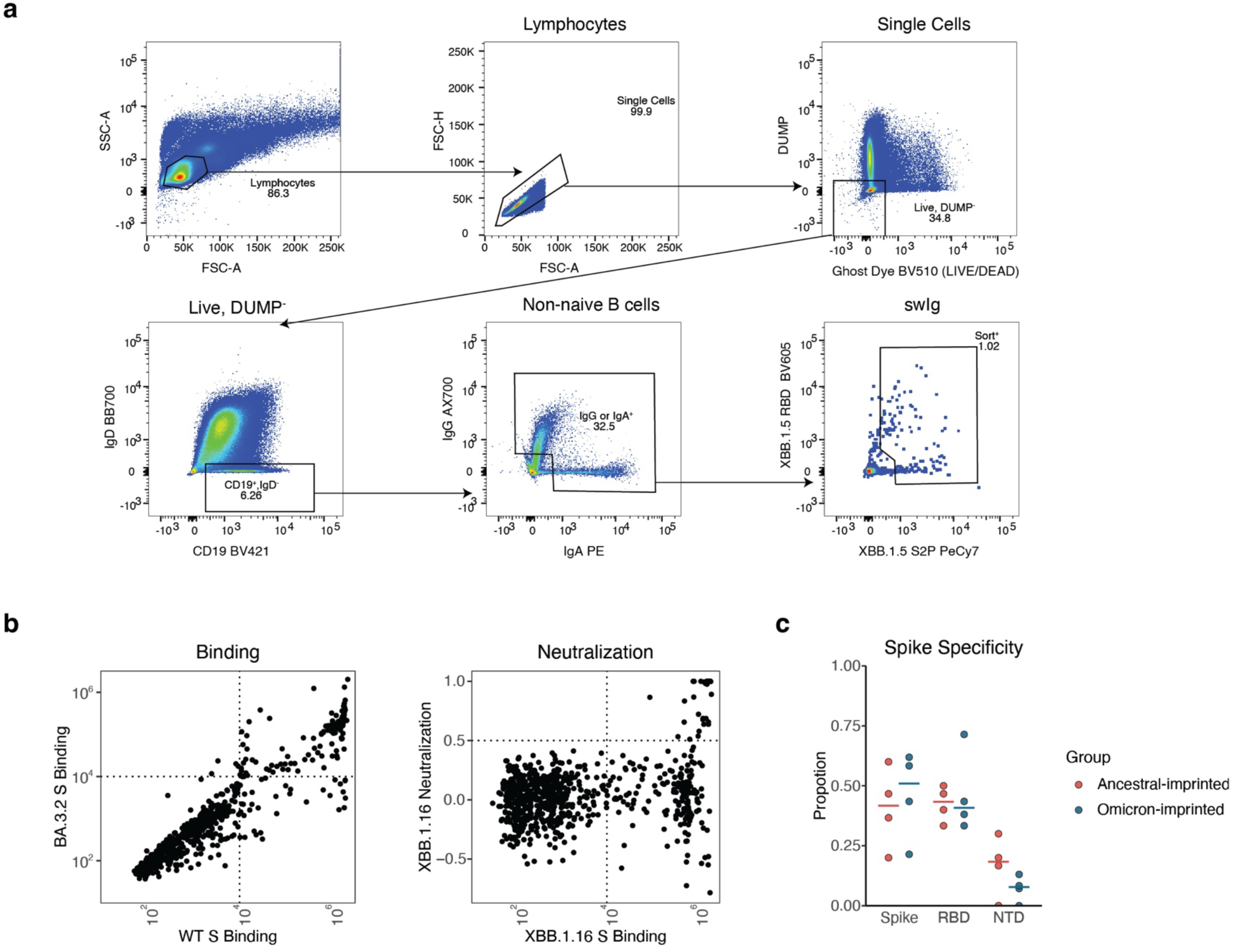
Isolation and analysis of monoclonal antibodies from children. a) Sorting scheme of class-switched memory B cells from peripheral blood mononuclear cells. b) Example plots showing a subset of variants for supernatant binding (left) and neutralization (right) screening. Each dot represents one monoclonal antibody supernatant. c) Proportion of monoclonal antibodies specific for full-length Spike, RBD, and NTD in each donor. Dots represent donors. Crossbars represent median.

**Extended Data Figure 5.**
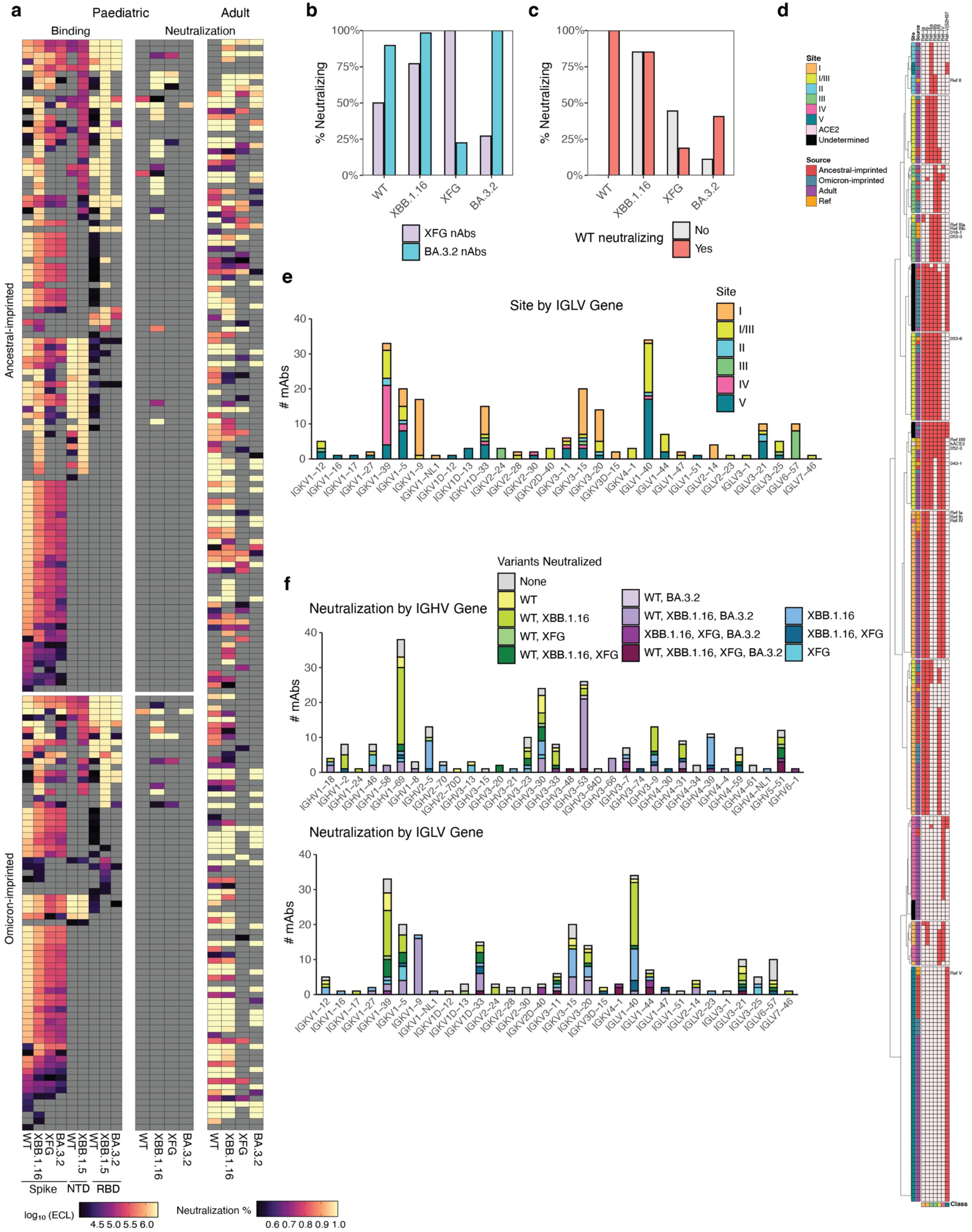
Monoclonal antibody analysis reveals differences in variant neutralization by RBD site. a) Heatmap of mAb binding and neutralization, with fill color denoting binding or neutralization intensity. b) Percent of XFG and BA.3.2 neutralizing antibodies that also neutralize WT, XBB.1.16, XFG, and BA.3.2. c) Percent of WT cross-reactive and Omicron type-specific neutralizing antibodies that neutralize WT, XBB.1.16, XFG, and BA.3.2. d) Epitope mapping clustering of unknown antibodies (y-axis) with reference antibodies (x-axis). Dark red = blocking, light red = binding. e) RBD binding site by IGHV gene usage. f) Variant neutralization by IGHV (top) and IGLV (bottom) mAb gene usage.

