## Supplementary Methods, Supplementary Table 1 & 2 for "Vaccine imprinting drives increased SARS-CoV-2 variant infection in children"

### Logistic regression – XFG and BA.3.2 sequence data

Let $I_i(t)$ be the density of hosts infected by the strain (or variant) $i$ of the pathogen at the current time $t$. The total density of infected hosts is then $I(t) = \sum_i I_i(t)$. Define $r_i(t)$, the growth rate of hosts infected by pathogen variant $i$. The temporal dynamics of $I_i(t)$ is given by the following ordinary differential equation:

$$\dot{I}_i(t) = r_i(t) I_i(t), \tag{1}$$

and the temporal dynamics of $I(t)$ by:

$$\dot{I}(t) = \sum_i r_i(t) I_i(t) = \underbrace{\sum_i q_i(t) r_i(t)}_{\bar{r}(t)} I(t), \tag{2}$$

where $q_i(t) = I_i(t)/I(t)$ represents the frequency of hosts infected by the variant $i$ among all infected hosts.

We are interested in tracking the frequency $q_i$:

$$\begin{aligned} \dot{q}_i(t) &= \frac{\dot{I}_i(t)}{I(t)} - q_i \frac{\dot{I}(t)}{I(t)} \\ &= q_i(t)\left(1 - q_i(t)\right) \underbrace{\left( r_i(t) - \frac{\sum_{j \neq i} q_j(t) r_j(t)}{\sum_{j \neq i} q_j(t)} \right)}_{\mathcal{S}_i(t)}, \end{aligned} \tag{3}$$

where we define $\mathcal{S}_i$ as the selection coefficient of variant $i$ relative to the background population, that is, the relative growth advantage (or disadvantage) of variant $i$ against all competing variants. Specifically, this coefficient measures the rate of change of the variant frequency on the logit scale – i.e., the log odds $\text{logit}(q_i/(1-q_i))$ –, and provides a relevant measure for the speed of pathogen adaptation:

$$\frac{\text{d logit}\left(q_i(t)\right)}{\text{d}t} = \mathcal{S}_i(t). \tag{4}$$

When $\mathcal{S}_i(t)$ is constant (or approximately constant), the variant frequency follows a logistic growth model: $\text{logit}\left(q_i(t)\right) = \text{logit}\left(q_i(0)\right) + \mathcal{S}_i \times t$.

We collected patient status metadata of GISAID SARS-CoV-2 sequences from New York and New Jersey US states (corresponding to the Health and Human Services (HHS) region 2, excluding Puerto Rico and the Virgin Islands) between 2025-01-05 to 2026-07-04. This region was chosen because it has a relatively large number of sequences compared to other locations, with good age coverage for those sequences. We consider all lineages associated in GISAID to Variant Under Monitoring (VUM) XFG + XFG.* as the XFG lineage, and all sequences associated to VUM BA.3.2 + BA.3.2.* as the BA.3.2 lineage.

We fit a linear generalized model (logistic regression) to sequence data. Let $Y_k(t) = 1$ if the $k$th sequence belongs to the variant $i$, and 0 otherwise, such that:

$$Y_k(t) \sim \text{Bernoulli}\left(q_i(t)\right). \tag{5}$$

We thus have:

$$\Pr\left[Y_k(t) = 1\right] = q_i(t), \tag{6}$$

assuming:

$$\text{logit}\left(q_i(t)\right) = \beta_0 + \beta_1 \times t, \tag{7}$$

with $\beta_0$, the intercept and $\beta_1$, the slope. The (relative) selection coefficient, $\mathcal{S}_i$, is thus estimated by the slope $\beta_1$. To minimize the influence of demographic stochasticity during the early phase of variant emergence, we initiate the logistic regression from the first day of the first week in which the variant frequency reaches 4%. We then restrict the fit of the model over the period during which the logit-transformed variant frequency is approximately linear. The final five weeks of data were excluded to account for delays in sequence reporting and backfilling.

We also fit a logistic regression to sequence data stratified by age groups:

$$\text{logit}\left(q_{i,a}(t)\right) = \beta_0 + \delta_a + (\beta_1 + \gamma_a) \times t, \tag{8}$$

with $\delta_a$, the difference in intercept for age group $a$ relative to the reference age group, and $\gamma_a$, the difference in slope. Here, the selection coefficient in age group $a$, $\mathcal{S}_{i,a}$, is estimated by the slope $(\beta_1 + \gamma_a)$. Here, we consider two age groups: individuals aged 0–10 years and individuals aged 11 years and older.

Table S1. Cohort Demographics

| | Adult | Paediatric | Ancestral (Paediatric subset) | Omicron (Paediatric subset) |
|---|---|---|---|---|
| **N** | 40 | 68 | 54 | 14 |
| **Age** | | | | |
| age min | 22 | 0.78 | 0.91 | 0.78 |
| age max | 59 | 3.96 | 3.96 | 3.59 |
| age median | 27.4 | 2.24 | 2.37 | 2.06 |
| **Sex** | | | | |
| M | 15 | 43 | 35 | 8 |
| F | 25 | 25 | 19 | 6 |
| **Race** | | | | |
| White | 28 | 56 | 44 | 12 |
| Black | 3 | 9 | 8 | 1 |
| Asian | 9 | 11 | 9 | 2 |
| **Ethnicity** | | | | |
| NHL | 39 | 55 | 47 | 8 |
| HL | 1 | 13 | 7 | 6 |

Table S2. Key Resources Table

| Reagent | Source | Catalog Number |
|---|---|---|
| **Antibodies** | | |
| BV421 anti-CD19 | Biolegend | Cat#302233 |
| BUV661 anti-CD38 | BD Biosciences | Cat#612969 |
| AF700 anti-IgG | BD Biosciences | Cat#561296 |
| FITC anti-CD3 | Biolegend | Cat#300406 |
| FITC anti-CD14 | Biolegend | Cat#325604 |
| FITC anti-CD16 | Biolegend | Cat#360716 |
| BB700 anti-IgD | BD Biosciences | Cat#566538 |
| PE anti-IgA | Miltenyi | Cat#130-116-878 |
| SULFO-TAG Anti-Human IgG Antibody | MSD | Cat#D21ADF-3 |
| **Chemicals, peptides and recombinant proteins** | | |
| SARS-CoV-2 Biotyinlated Full Length Spike (WT, XBB.1.16, XFG, BA.3.2) | In-house | |
| SARS-CoV-2 Biotyinlated Receptor Binding Domain (WT, XBB.1.5, BA.3.2) | In-house | |
| SARS-CoV-2 Biotyinlated N-Terminal Domain (WT, XBB.1.5) | In-house | |
| BV605 Streptavidin | Biolegend | Cat#405229 |
| PE-Cy7 Streptavidin | Biolegend | Cat#405206 |
| Ghost Dye 510 Viability Dye | Tonbo | Cat#13-0870-T100 |
